# CRISPR-VAE: A Method for Explaining CRISPR/Cas12a Predictions, and an Efficiency-aware gRNA Sequence Generator

**DOI:** 10.1101/2021.07.05.451176

**Authors:** Ahmad Obeid, Hasan AlMarzouqi

## Abstract

Deep learning has shown great promise in the prediction of the gRNA efficiency, which helps optimize the engineered gRNAs, and thus has greatly improved the usage of CRISPR-Cas systems in genome editing. However, the black box prediction of deep learning methods does not provide adequate explanation to the factors that increase efficiency; rectifying this issue promotes the usage of CRISPR-Cas systems in numerous domains. We put forward a framework for interpreting gRNA efficiency prediction, dubbed CRISPR-VAE, that improves understanding the factors that increase gRNA efficiency, and apply it to CRISPR/Cas12a (formally known as CRISPR/Cpf1). We further lay out a semantic articulation of such factors into position-wise k-mer rules. The paradigm consists of building an efficiency-aware gRNA sequence generator trained on available real data, and using it to generate a large amount of synthetic sequences with favorable traits, upon which the explanation of the gRNA prediction is based. CRISPR-VAE can further be used as a standalone sequence generator, where the user has low-level control ability. The framework can be readily integrated with different CRISPR-Cas tools and datasets, and its efficacy is confirmed. The complete implementation of the methods can be found at github.com/AhmadObeid/CRISPR-VAE.

## 1 Introduction

THE usage of CRISPR-Cas systems for genome editing has been gaining much popularity recently due to the many applications which the technology enables in various domains such as gene therapy and agricultural engineering [1], [2], [3], [4]. Such popularity motivated an advancement in the related research, particularly including many works towards the prediction of guide RNAs (gRNAs) efficiency. In CRSIPR-Cas systems, gRNAs locate DNA targets for the endonuclease to cleave. When the DNA strands are being repaired, the process results in random insertions/deletions or precise gene editing that can be exploited for gene knockins [5]. The efficacy of said process is thus a function of the used gRNA; predicting it is important for a safe usage of CRISPR-Cas systems, in order to ensure high on-target indel efficacy, and minimum off-target effects. Additionally, the discovery of Cas12a, also known as CRISPR from Prevotella and Francisella1 (Cpf1) as an alternative endonuclease to CRISPR associated protein 9 (Cas9) in the CRISPR systems is an important development, and introduces many favorable features. For example, Cpf1 is shorter in size, requires a smaller CRISPR Ribonucleic Acid (RNA) to function, and facilitates the re-engineering of the desired DNA as the target, with the Protospacer Adjacent Motif (PAM) remaining unaffected [6]. Additionally, Cpf1 has shown better specificity in human and plant cells than Cas9, and enables the editing of Corynebacterium glutamicum and Cyanobacteria, which was not possible with Cas9 [5].

The methods used for gRNA activity prediction can be alignment-based, hypothesis-driven, or learning-based [7]. Alignment-based methods rely entirely on locating the PAM, in order to align the gRNA in the genome. Hypothesis-driven methods score the aligned gRNAs by other contextual factors. Learning-based methods train a prediction model that can extract many hidden sequence-related factors to infer the on-target efficiency of the sequence. With the continuous advancement in the area of deep learning, learning-based methods have been showing high accuracy and a promising performance at gRNA efficiency prediction. However, these methods are still inadequately interpreted, and provide little explainability to their predictions. Said explainability is crucial for a better understanding of CRISPR systems, and to explore the factors that make certain gRNAs lead to higher on-target activities. This enables practitioners design better sequences, and analysts better diagnose their models’ decisions, which promotes the application of genome editing in different domains.

Previous attempts have touched upon this research direction [5], [7], [8], [9], but were faced with the two challenges of deficient and incomprehensive data. The deficiency in the data is represented by having relatively few sequences that belong to specific efficiency categories. This, in turn, disallows the establishment of statistically significant factors. As for the incomprehensiveness of the data, we are referring to the case where available datasets consist of an incohesive collection of sequences that exhibit many sequence-related features, where no clear connection can be drawn. This scatters the effort of finding the features that are responsible for high editing efficiency. In this work, we develop a framework that tackles both problems simultaneously, as illustrated in Fig.1. More concretely, publicly available data of CRISPR/Cpf1 activity leaves sequence and structure-related gaps in a certain analysis space (to be explained later), which we place a magnifying lens over; manifested in the development of a sequence generator dubbed CRISPR-Variational Autoencoder (CRISPR-VAE).

**Fig. 1.**
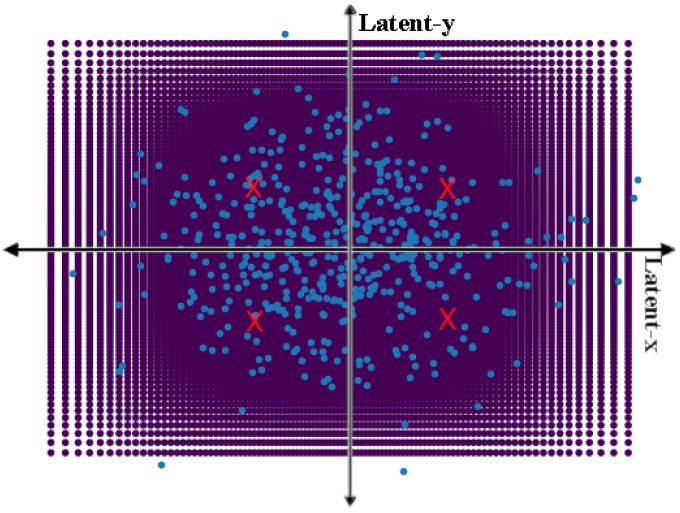
The latent space obtained by CRISPR-VAE, showing the seeds of the synthetic data (dark and regular grid) bridging sequence-related gaps left by publicly available data (light and scattered points). The cross-marks indicate the quadrant centroid

CRISPR-VAE is efficiency-aware, and is used to synthesize numerous sequences of high and low efficiencies. These sequences are not arbitrary, but form a structured analysis space that is meant to bridge said gaps left by the dataset, and exhibits different sequence phenomena ordered in different locations in the space. As such, this produces more comprehensive and plentiful data, upon which the discovery of rules is based. Such a paradigm concentrates and magnifies the search for efficiency-promoting factors. Finally, we predict the efficiency of the synthetic sequences using the deep learning-based predictor seq-DeepCpf1 which previously showed good performance on the dataset [10]. As such, we establish an agreement between two methodologically distinct frameworks: a generative one and discriminative one, which increases the confidence in the findings.

In summary, the contribution of our work is:

- putting forward a deep learning-based explainability framework, which can be readily integrated with any CRISPR-Cas dataset;
- developing methods that improve the statistical significance of findings, and concentrates the search for them;
- semantically articulating the high-quality findings of the mentioned methods in a suitable k-mer paradigm;
- developing the standalone sequence generator CRISPR-VAE, which can generate sequences of high (or low) efficiencies with low-level, position-wise features tailored to the practitioner’s needs;
- demonstrating the correctness of findings by establishing an agreement between a descriminative and a generative method.

In the following, we will provide a literature review in Section 2 for the related works in the themes of deep learning-based prediction of gRNA efficiency and explainability. In Section 3, we will describe how a structured analysis space is obtained, how an efficiency-aware gRNA generator is made, and how the different sequence features are extracted. Finally, we summarize the results of our experimentation, and mention some concluding remarks in Sections 4 and 5.

## 2 Related Work

The task of predicting a quantifiable quality assessment of a gRNA sequence is realized through predicting its on-target efficiency. In this vain, the ability to flesh-out well-articulated rules that can be interpreted by humans further helps practitioners decipher the genome code. Consequently, researchers have been developing analytical tools for the mentioned tasks, among which deep learning-based tools have been showing special promise due to the continuous advancement in the field [7], [10], [11], [12], [13], [14]. In spite of that, the issue of interpretation in deep learning-based tools towards the prediction of gRNA efficiency is still immature, and such methods still lack strong explainability that can guide the analysis in a meaningful fault-diagnosis direction.

In [10], the DeepCpf1 and seq-DeepCpf1 predictors have been developed using Convolutional Neural Networks (CNNs) and dense ones. Said predictors show an improved performance in comparison to other classical methods. In [11], the authors follow a similar path, but the support vector regression (SVR) is used at the end to aid the CNN network, showing some improvements in the performance. Nonetheless, both proposed paradigms, which target the Cpf1 endonuclease, do not provide any explainability for their prediction. Similarly, the DeepCas9 [13] makes use of CNNs, DeepCRISPR [12] aids them with an Autoencoder (AE) stage for unsupervised representation learning, and the C-RNNCrispr [7] aids them with a Recurrent Neural Network (RNN) for a better sequence learning. All these methods and others have been developed for the cas9 endonuclease. Moreover, DeepPE [14] was introduced for the new tool of Prime Editing, also making use of CNNs. Despite the versatility and prowess introduced to tackle the task of gRNA efficiency prediction, most of such paradigms do not tackle the issue of explainability, and deal with their predictors as black boxes. Additionally, there are various works that employ deep learning for gRNA off-target prediction [15], [16], [17], also suffering from the black-box behaviour of their predictors.

On the other hand, the issue of explainability in deep learning-based prediction of gRNA efficiency has been studied. For example, [7] studies an optimization of their model score with respect to the inputted gRNA sequence, in order to infer the most prominent features that maximize the efficiency. A similar approach is also used in [18]. Contrariwise, some works opt for classical machine learning tools that are easier to explain [19], thus trading-off accuracy with explainablity.

Other attempts that employ statistical analysis of the available data to infer position-wise base preference rules [5], [8], [9] are relevant, although they do not employ deep learning-based methods for their rule inference. However, the main issue with such methods is that they base the analysis exclusively on the available data, which exhibits a few limitations. Firstly, the available data is limited in quantity (i.e. class-wise), despite publishing large datasets for both cas9 and Cpf1 endonucleases. For example, the available data on Cpf1 has a small number of sequences with efficiency ≥ 0.99 or ≤ 0.05, although having a large number of such greatly polarized sequences is important to infer the most prominent rules to a statistically significant degree [20], [21]. More gravely, the available data suffers from qualitative limitation. In other words, the available data is usually scattered in a sequence-cohesion sense, exhibiting different and distinct sequence-related features. Each one of these features is also obscurely represented in the data, making it difficult to discover them. Theoretically, a quantitative comprehensiveness cannot be realistically achieved. Instead, we seek to meaningfully expand the data to signify the different features both quantitatively and qualitatively. In this work, we focus on building a framework that is well embedded in the deep learning paradigm, and that can be generically applied to any CRISPR-Cas system, while tackling the mentioned difficulties in the available data.

## 3 Materials and Methods

In this Section, we will describe the three main components of our proposed framework, which starts with the proposed generative framework CRISPR-VAE and its advantages, then describes the subsequent feature extraction procedure.

### 3.1 VAE for a Structured Latent Space

We start by describing the general paradigm of VAE, which enables establishing a structured latent space, and the benefit of the latter.

The illustration in Fig.1 demonstrates the core of the analysis paradigm. In contrast to previous attempts, the proposed work aims to accentuate and distinguish the existing sequence-related phenomena, and explore possible ignored ones in their neighborhood. This necessitates establishing a structured analysis space, for which we employ the VAE paradigm. The analysis space is composed of numerous synthetic sequences, that share a resemblance with the training data, and that are systematically distributed over the space.

The generative process of the VAE starts by generating latent variable **z** from the prior distribution *p*_*θ*_(**z**). Then, **x** is generated (reconstructed) from the generative distribution *p*_*θ*_(**x**|**z**). In this framework, parameter estimation is difficult due to the intractability of the posterior. Alternatively, the lower-bound of the log likelihood is used:

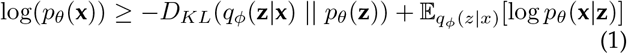

where *q*_*ϕ*_(**z**|**x**) is an approximation for the true posterior *p*_*θ*_(**z**|**x**), and *D*_*KL*_(. || .) is the KL-divergence. In our implementation, we use CNNs and dense layers for the realization of both models *p*_*θ*_(**x**|**z**) and *q*_*ϕ*_(**z**|**x**) i.e. the encoder and decoder models, respectively, as shown in Fig. 2. Furthermore, assuming a Gaussian latent variable, the empirical loss of the VAE can be written as:

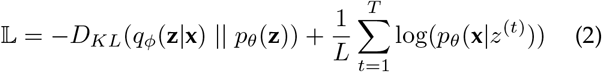

where *z*^(*t*)^ is a sample drawn from the generative model i.e. *z*^(*t*)^ = *g*_*ϕ*_(**x**, *ϵ*), and *ϵ* ∼ 𝒩(0, **1**) is used for the so-called reparametrization trick [22].

**Fig. 2.**
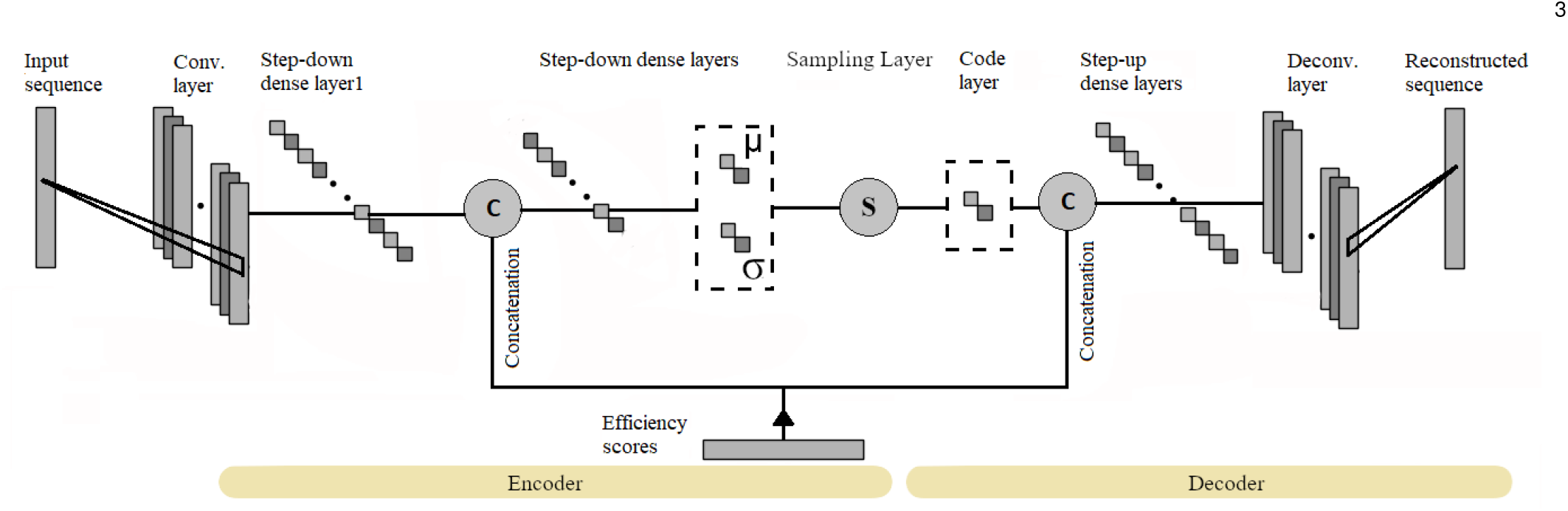
CRISPR-VAE architecture, shown to integrate the efficiency information at two stages: end of encoder, and beginning of decoder

The encoder model learns to project the sequences into the latent space, assimilating them into a normal distribution, where different sequence-related features occupy different locations of the space. Nevertheless, the projected sequences of the training data exhibit a non-cohesive latent space, leaving certain gaps for exploration. As such, a systematic and structured sampling from the latent space for decoding will result in the synthesis of novel sequences that resemble the original data, and that fill in the left gaps, which provides a continuous and smooth bridging between the different sequence phenomena.

The shown latent space in Fig. 1 is two-dimensional (2D), but the analysis can be extended to higher dimensions, giving more prowess to the VAE architecture, and thus obtaining a higher quality of reconstruction and synthesis, and a larger amount of synthetic data. This comes over the expense of a more complex analysis and a demand of higher storage capacity.

The benefit of having a structured latent space is two-fold. Firstly, we ensure that all phenomena existing and scattered in the original data are parsed and highlighted. Secondly, it was empirically demonstrated that the structured space systematically distributes the different sequence phenomena in its different locations (e.g., quadrants in 2D space). This eases the analysis and search for efficiency-promoting features. Additionally, this gives the sequence generator a low-level capability of editing, where specific position-wise base preferences are translated to sampling from different quadrants.

Moreover, without a structured latent space for analysis, exploring ignored potential phenomena in a comprehensive manner would be too wasteful and demanding of resources, with an estimated upper-limit complexity in the search space of 𝒪(31^4^) (assuming gRNA sequences of length 34, with a known PAM of TTTV), where the vast majority of such sequences would have no connection to the available data, rendering the validation impossible. Instead, the proposed framework enables the synthesis to be confined to sequences that resemble the available data. In our implementation, we sampled 10, 000 latent codes arranged in a grid of 100×100 which are decoded to synthetic sequences for subsequent analysis stages.

To test the structure of the latent space, distance heat maps have been constructed through the following:

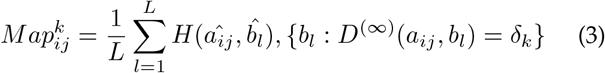

where i and j denote each point in the heat map, k denotes the specific heat map that corresponds to the used 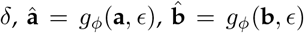, *H* is the Hamming distance, and *D*^(∞)^ is the Minkowski distance of order ∞. L denotes the number of seeds in the latent space that satisfy the Min-skowski distance condition. A structured space is expected to exhibit heat maps with values growing proportional to *δ*.

### 3.2 CVAE for efficiency-awareness

In our implementation, we specifically follow the conditional VAE (CVAE) paradigm inspired by [22], where we condition on the efficiency score of each sequence. In this Section, we will describe the needed change that grants CRISPR-VAE its efficiency-awareness.

More concretely, equation (2) becomes:

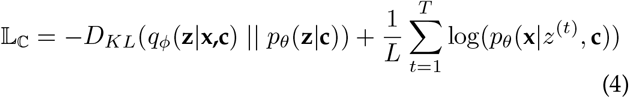

where we are conditioning the encoding, the decoding, and the prior distribution on the efficiency information *c*. This means that we obtain a separate latent space for each efficiency category. In our implementation, we convert the efficiency scores in the public data to integers, resulting in a 100 classes (0 − 99). Theoretically, this results in up to 100 latent spaces, each consisting of 100×100 grid of sequences. However, we confined the synthesis to classes 0-efficiency and 99-efficiency to synthesize the most polarized sequences, in order to focus on the most prominent and distinct sequence-related features that set the high-efficiency sequences apart from their low-efficiency counterparts.

Integrating the CVAE paradigm has two benefits. Firstly, the available data provides efficiency measurements which represent useful information to improve the quality of synthesis and reconstruction of the VAE. Indeed, experimentation showed that exploiting the efficiency information improves the performance of the VAE, as will be shown in Section 4. Secondly, exploiting the efficiency information of the data makes the VAE efficiency-aware, and makes the synthesis of the data tailored to the needs of the user (e.g., focused on the high-efficiency sequences). Moreover, this enables us to prob the agreement between the CVAE and existing descriminative methods. In Section 4, we show that CRISPR-VAE and the seq-DeepCpf1 predictor [10] have a strong agreement, and thus increasing the confidence in the findings, without the need for laboratory testing. Herein, another benefit of having a structured latent space is apparent, where we can enforce a resemblance between the synthetic data and the original data, making seq-DeepCpf1 familiar with the synthetic data.

A final trick was employed to improve the quality of reconstruction of CRISPR-VAE. Particularly, the first three loci in the PAM of all sequences were removed, as they are constantly TTT. It was observed upon experimentation that the model finds reconstructing such motif as an easy way to score highly in the objective function; avoiding it motivated the model to rely on learning more interesting sequence-related features that improve the quality of reconstruction, which had a direct impact on said quality.

Fig.2 illustrates the CVAE paradigm. The one-hot-encoded efficiency information is fed to the network at two concatenation stages. The first stage, which comes after the first dense layer, allows the efficiency information to be blended and integrated with the sequence information via the subsequent dense layers, and into the embedding of the code layer, establishing the latent space. The second stage, which comes after the code layer, allows the decoder to be a standalone efficiency-aware sequence generator. The sampling layer employs the aforementioned reparametrization trick to convert *µ* and *σ* to the latent codes.

### 3.3 Data usage

As efficiency scores are used, it is applicable to split our data into training and testing sets to confirm the generality of our findings. We use the data provided in [10], where high-throughput experiments were used to generate sets HT1, HT2, and HT3. We use set HT1 for training, which consists of ∼16300 sequences, while sets HT2 and HT3 were used for testing. These sets do not share any sequences, which excludes any possibility of data leakage. We also applied data augmentation by causing small perturbations in the promiscuous region of each sequence in HT1 such that the efficiency scores are likely maintained according to [9], resulting in *∼*85,000 sequences to train CRISPR-VAE.

### 3.4 Feature Extraction

Having built CRISPR-VAE, one can synthesize numerous sequences that exhibit the two main characteristics missing from the original data: comprehensiveness and plentifulness. What remains is to extract the sequnce-related features that are responsible for the disparity in the sequences that belong to 0-efficiency and 99-efficiency classes. To that end, two methods were used to extract and articulate the most prominent features explored in the synthetic data. The first one consists of k-mer histogramming analysis to build Mer Significance Maps (MSMs), and the second consists of visualizing class activation maps (CAMs) [23] produced by a binary classifier that distinguishes the high-efficiency from the low-efficiency sequences.

Firstly, following [10], we define the three regions of seed, trunk, and promiscuous, in addition to PAM, pre-PAM, and post-seq regions, as shown in Fig. 3. In the first method, we employ a moving overlapping window encapsulating k-mers to segregate the feature extraction based on the position in the gRNA sequence. This confers the analysis with the needed contextual specificity. We chose the step size to be 1 base, resulting in *L* − *k* + 1 sub-regions for each parent region, where *L* is the length of the region (e.g., *L*_*trunk*_ = 12), and *k* is the mer size. In this paper, we constricted the experimentation to *k* = 3, but other options can be easily realized. Also, we split the latent space into equal-sized locations (e.g., 4 quadrants in 2D space), where the histogramming takes place independently. This is to highlight certain phenomena that may otherwise be overshadowed by more prominent ones. It was indeed observed that different sequence features are prominent in different locations in the latent space.

**Fig. 3.**
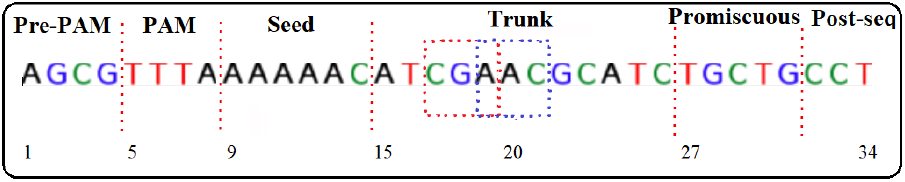
Different regions in gRNA, with an example of two overlapping mer windows of size 3 in the trunk region

After pooling in all possible features, we filtered them by firstly setting an empirical significance threshold (*η*) where features with frequency below this threshold are discarded of. The threshold is chosen as *m*% (typically 7∼10%) of the number of possible mers, multiplied by the number of sequences under analysis (*N*), as shown in equation (5). *N* can refer to the whole set of generated sequences, or only sequences in a specific quadrant. *η* is defined for each quadrant, for each region in the gRNA, and for high-efficiency and low-efficiency sequences, separately, albeit with a constant *m*% throughout. Secondly, we discarded of features that are simultaneously above the low-efficiency and high-efficiency significance thresholds i.e. the common features between the two classes. Moreover, we highlight some novel features that are obscure in the training data, but are discovered due to the mentioned benefits introduced by the synthetic data, and are later found to exist in the testing data. In other words, these features would have likely been ignored if the analysis did not involve synthesizing sequences (i.e. similar to [5]), alluding to the added generality conferred by the proposed framework. We can thus point out the differences between the proposed paradigm and that of [5] as introducing more specificity by exploiting the hidden cohesion in the training data, more sensitivity by highlighting possibly overshadowed features, and more generality by discovering the neighborhood of the real sequences.

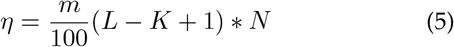

In the second method, we trained a binary classifier on the synthetic data, and visualized the CAMs [23] for each quadrant separately. Said CAMs are obtained by maximizing the score of the binary efficiency classifier with respect to the first convolutional layer. This results in attention maps that visually describe the reason for the decision of the classifier. We obtained such maps for the decision of all 99-efficiency sequences, and averaged them. Such features can be easily compared with their counterparts from the first method, thus enabling testing the agreement between both methods. Moreover, the two methods are complementary to each other. The first method has a finer granularity in terms of the location of the prominent features, while the second is an automatic method that directly explains the decision of the binary classifier. By looking at the results of both methods, numerous features can be extracted, and a better understanding of the composition of an efficient sequence can be made. Finally, we use Welch’s t-test to evaluate the statistical significance of the explored features. As such, we include the p-values of observing a high-efficiency sequence, exhibiting each of the explored features.

The full code of the methodology, including all filtering stages, and all generated results is shared publicly.^1^

## 4 Results

We split the Results Section into two parts. In the first one, we show results that confirm the efficacy of the proposed paradigm. Secondly, we summarize the inferred sequence-related features from the two methods mentioned in 3.4. Moreover, in Section S.I of the supplementary material, we include an explanation of how CRISPR-VAE can be used as a sequence generator, and the method by which users can control the nucleotide content of their synthetic sequences from the 99-efficiency class.

### 4.1 Confirming the Validity of the Proposed Framework

We start by confirming that the constructed latent space is structured. A structured latent space exhibits smooth transitioning between the different sequences, placing similar sequences in each others’ vicinity. As such, when varying *δ* (equation (3), a structured latent space should give heat maps with values growing proportional to it. In Fig. 4, *δ* = {3, 11, 29} in order to compare sequences with small, medium and large separations. The shown heat maps are for the class 99-efficiency sequences. Indeed, the heat maps behave as expected from a structured space, with the smallest values observed with *δ* = 3, and the largest with *δ* = 29.

**Fig. 4.**
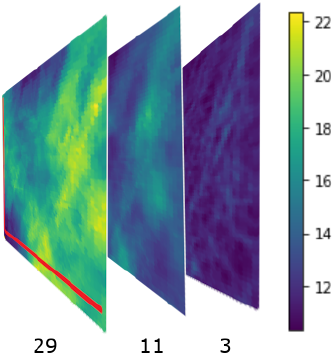
Results of heat maps generation by equation (3), showing a monotonic positive relation between *δ* = 3, 11, 29 and the Hamming distance, clearly indicating the sequential structure in the synthetic latent space.

Furthermore, Fig. 5 provides a pictorial agreement assessment between the generative CRISPR-VAE and the descriminative seq-deepCpf1 predictor. The figure shows the prediction of the seq-deepCpf1 method on synthetic data claimed to belong to classes 0-efficiency and 99-efficiency by CRISPR-VAE. Ideally, the figure should exhibit two disjoint peaks located at the two extremes of 0 and 99. The figure indeed shows a tendency towards such behaviour, which can be quantified by a Spearman’s correlation coefficient of *∼* 0.823. In other words, according to seq-deepCpf1, most sequences claimed for class 99-efficiency have higher efficiencies than most sequences claimed for class 0-efficiency, which enables the identification of the most prominent features that set the two types of sequences apart.

**Fig. 5.**
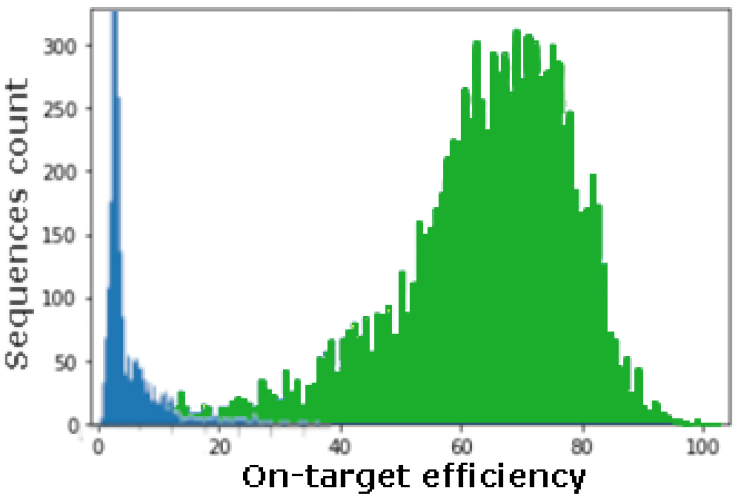
seq-deepCpf1 [10] prediction on the synthetic data of efficiencies 0 (left) & 99 (right)

### 4.2 High-efficiency Features

Herein, we include Fig. 6 for a holistic summary of the sequence-related features in different sequence regions and quadrants in the latent space. Said features pass the filtering stages and are significantly prominent in the sequences of class 99-efficiency in contrast to sequences of class 0-efficiency. Filtering with the significance threshold results in focusing on the prominent features, and consequently having some empty sub-regions in Fig. (6,c). The MSMs consist of cocentric significance circles, whose radii are proportional to their significance. The different regions are color-coded as per the legend in Fig. (6,a). The discovered mers are scattered in the MSMs based on their significance and position in the gRNA; the larger the angle at which the mer is located, the more down the gRNA stream the mer feature exists. For clarity, we also segregated each region into three sub-regions separated by dash lines. These sub-regions describe the beginning, the middle, and the end of each region. In Fig. (6,c), we highlight the significant features found in the synthetic data which are obscure in the training data HT1, but are confirmed by the testing data HT2 and HT3 by circling them.

**Fig. 6.**
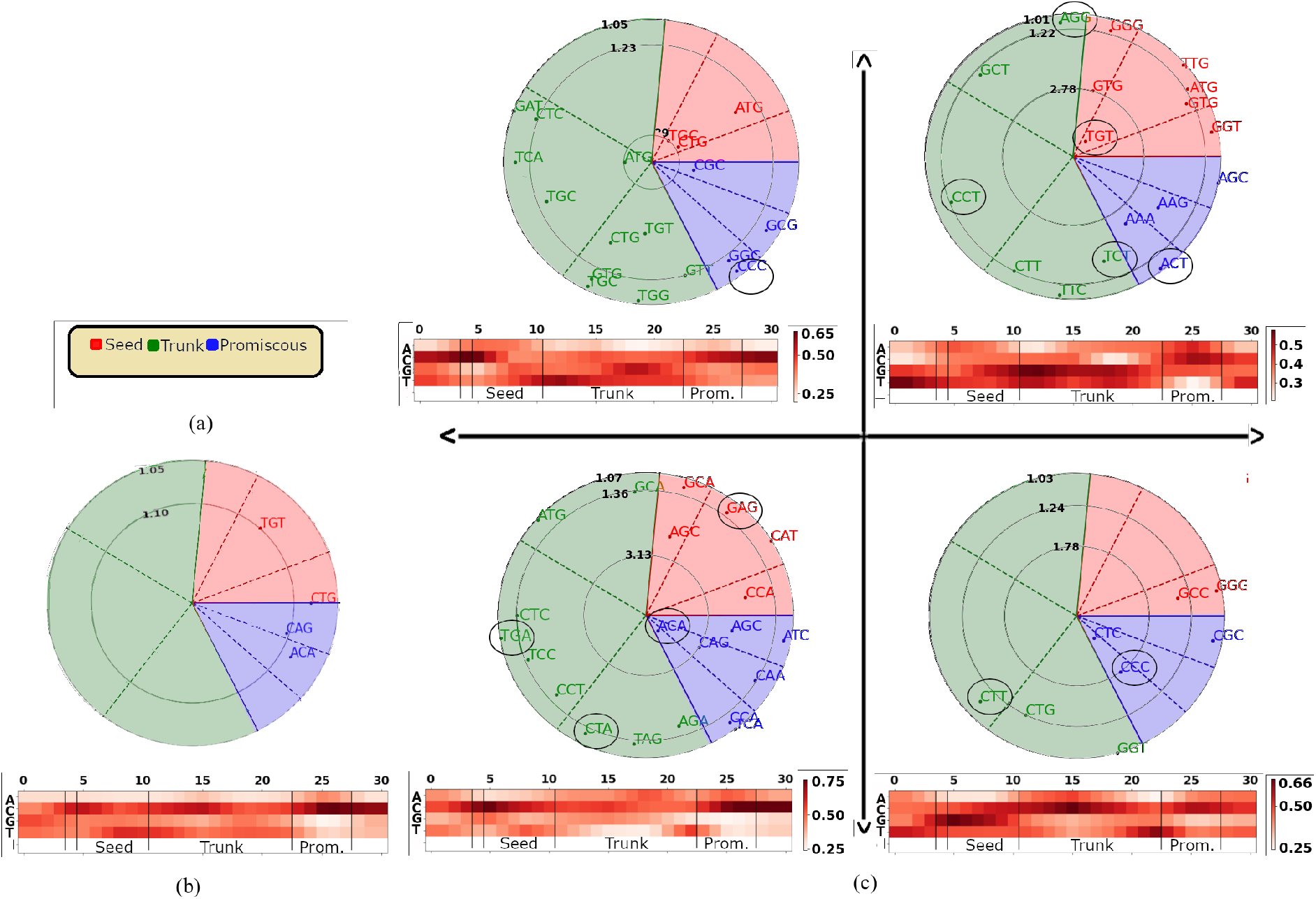
Summary of high-efficiency sequence-related features as MSMs and CAMs, showing the benefit of a quadrant-based analysis (c), as compared to non-quadrant-based (b). The sequence regions are color-coded according to the legend in (a), and separated into three sub-regions (beginning⟶mid⟶end) going counter-clockwise; the circled mers in (c) can be exclusively found from the synthetic data, and not the training data, alluding to the added generality conferred by the proposed paradigm. In the MSMs, more significant features are closer to the center. Only the last nucleotide in the PAM is shown in the CAMs.

Fig. 6 also provides a pictorial summary of the distinguishing trends that set the two types of sequences apart using CAM via a specialized classifier. The classifier learns to classify the synthetic data to a near-perfect degree with only few epochs (accuracy ∼95%, with 5 epochs), alluding to how distinct the features in both categories are. We also include Fig. (6,b) to show the benefit of segregating the analysis to different quadrants in the latent space. Otherwise, the analysis reveals an averaged version where only the globally prominent features are highlighted while ignoring many other valid ones. For example, the averaged summary in Fig. (6,b) reveals a disfavoring of Thymine right after the PAM, which agrees with the existing findings [5], but does not reveal much more than that.

Many sequence-related features that result in high-efficiency sequences can be inferred by looking at Fig. 6. We focus especially on the features that are agreed upon by the two methods of CAMs and MSMs. Firstly, Adenine is preferred mainly in the promiscuous region, as shown in quadrants1 and 3 (Q1 and Q3), as shown in the CAMs, and confirmed by some mers in the MSMs. Adenine therein especially prefers to combine with Cytosine or Guanine in said region (e.g., AAA and AAG in Q1, ACA and CAA in Q3).

As for Cytosine, it can be positioned everywhere in the gRNA, albeit in different combinations depending on the region, as revealed looking at the different quadrants. More concretely, for Cytosine in to be in the seed region, it prefers to combine with the TG pair (e.g., TGC and CTG in Q2), or preceded by Guanine (e.g., AGC, GCA in Q3), or followed by Adenine (e.g., GCA, CCA, CAT in Q3). As such, one can conclude that a motif of TGCA is observed in the seed region of efficient sequences. For Cytosine to be in the middle towards the end of the trunk region, it prefers to be combined with Thymine as observed in different mers in various quadrants. As for the promiscuous region, Cytosine prefers to either combine with Guanine (e.g., CGC, GCG in Q2) or be followed by Adenine (e.g., ACA, CAG, CAA, CCA, TCA in Q3).

As for Guanine, it can be placed at the beginning of the seed region if followed by Cytosine as revealed in Q4 (e.g., GCC), or anywhere in the seed region if preceded by Thymine or Adenine (e.g., TGT, TTG, GTG, ATG), or in the beginning of the trunk region (e.g., AGG), as revealed in Q1.

Thymine makes an appearance in various places. It prefers to combine with Cytosine in the middle towards the end of the trunk region as revealed in Q1, Q3, and Q4. For Thymine to be in the beginning towards the middle of the trunk region, it prefers to be followed by Guanine (e.g., TGT, TGG, TGC, GTG) as shown in Q2. Thymine is disfavored in the seed and promiscuous regions, except as auxiliary bases in a few cases.

The aforementioned features and many others can be inferred and summarized, particularly those that have been revealed exclusively by the synthetic data. In the MSMs, multiple such features are included, such as the TGT mer in the seed region, ACT, CCC, and ACA in the promiscuous region, and many other ones in the trunk region across the different quadrants. These features exist in the testing data HT2 and HT3, but are obscurely observed in the training data HT1. This showcases the direct benefit of the suggested paradigm, where it is possible to discover obscure features that lie in the neighborhood of the prominent ones of the training data. For results of testing the statistical significance of the mers using Welch’s t-test, the reader is referred to Section S.II of the supplementary file. Moreover, for an explanation of how CRISPR-VAE can be used as a sequence generator with low-level editing control, Section S.I of the supplementary document is referred to.

## 5 Conclusions

In this paper, we developed a complete paradigm towards improving the explainability of deep learning-based models in the application of gRNA sequence efficiency prediction in CRISPR systems. The paradigm consists of building a generative framework where synthetic data is generated that resembles labeled training data, and fills in the sequence-related gaps in it. The agreement between the proposed generative framework and the descriminative seq-DeepCpf1 increases the confidence in the findings, and also provides explainability for the decision of the descriminative method. Two analysis methods were used to infer and summarize the most prominent features from the synthetic data. The first is a manual histogramming method, and the second is automatic, using class activation maps. Many features have been thus discovered and highlighted, including particularly obscure ones that are confirmed to be in the testing data. Lastly, we showcased in the supplementary document the capability of the proposed framework in generating gRNA sequences, with a low-level editing control, by altering the latent code. We further mapped out the relationship between the position of the latent code and the expected features in the generated sequence.

## Acknowledgments

This work was supported by the Abu Dhabi Award for Research Excellence under ASPIRE/Advanced Technology Research Council

---

**Ahmad Obeid** received the B.S. degree in Electrical Engineering from the American University of Sharjah in 2018, and the M.S. degree in Electrical Engineering and Computer Science from Khalifa University, Abu Dhabi, in 2020 where he is also currently pursuing his Ph.D. studies in Electrical Engineering and Computer Science.

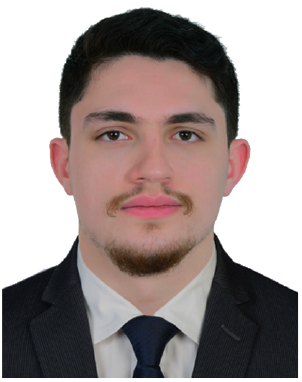

After receiving his M.S. and before commencing his Ph.D studies, he served as a Research Associate at Khalifa University. He is the author of several papers in conferences and journals, and has been involved in versatile research projects. His research interests include signal processing, machine learning, and computer vision.

Mr. Ahmad was a recipient of several research awards and scholar-ships, including the Sharjah Chamber of Commerce and Industry Award for Innovators. He is also a member Eta Kappa Knu and Tau Beta Pi Honor societies.

**Hasan Al-Marzouqi** (Senior Member, IEEE) received the bachelor’s (Hons.) and M.S. degrees in electrical and computer engineering from Vanderbilt University, Nashville, TN, USA, in 2004 and 2006, respectively, and the Ph.D. degree in electrical and computer engineering from the Georgia Institute of Technology, in 2014. He is currently an Assistant Professor with the Electrical and Computer Engineering Department, Khalifa University, Abu Dhabi, United Arab Emirates. His research interests include image and video processing, machine learning, bioinformatics, and digital rock physics. He served as an Organizing Committee Member and the Challenge Sessions Co-Chair of ICIP 2020 in Abu Dhabi and a Technical Program Committee Member of EUVIP 2020 in Paris. He is an active reviewer of many international conferences and journals.

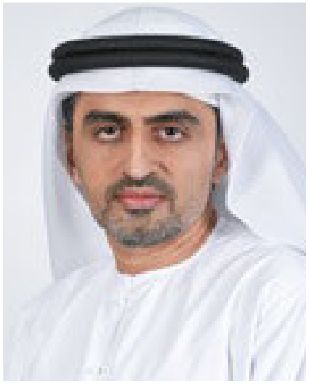

## Footnotes

1 github.com/AhmadObeid/CRISPR-VAE

